# Laccase-mediated biotransformation potential for fluorinated compounds by geographically diverse human gut microbiota

**DOI:** 10.64898/2026.04.27.720343

**Authors:** Milo R. Schärer, Yaochun Yu, Aaron Grawe, James K. Christenson, Serina L. Robinson, Nicholas A. Bokulich

**Affiliations:** Institute of Food, Nutrition and Health, Department of Health Sciences and Technology, ETH Zurich, Zurich, Zurich, Switzerland; Department of Environmental Microbiology, Swiss Federal Institute of Aquatic Science and Technology, Dübendorf, Zurich, Switzerland; Department of Chemistry and Biochemistry, Bethel University, St. Paul, Minnesota, United States of America; Institute of Biogeochemistry and Pollutant Dynamics, Department of Environmental Systems Science, ETH Zurich, Zurich, Zurich, Switzerland

**Author notes:** Department of Civil and Environmental Engineering, Stanford University, Stanford, California 94305, United States of America (Yaochun Yu). co-first (Milo R. Schärer, Yaochun Yu).

## Abstract

The growing prevalence of synthetic organofluorine substances in agrochemicals, food packaging, and consumer products has led to increasing gastrointestinal exposure, with potential consequences for human health. Despite the extreme stability of fluorinated compounds, several microbial pathways for their transformation are known, including by laccases, a type of multicopper oxidase. However, the functionality of laccases in the gut microbiome, a natural contact point between food-associated chemicals and microbial biotransformation pathways, is poorly defined. Through a multi-study analysis of 1578 human gut metagenomes spanning a global gradient from hunter-gatherer societies to industrialized urban populations, we found that laccase-coding gene homologs are widely distributed in the human gut microbiome. We identified a significant association between both the abundance and phylogenetic diversity of laccase homologs with the degree of urbanization. As human gut microbial laccase activity has not been experimentally demonstrated, eight gut metagenome-derived laccases were heterologously expressed and screened for activity with a redox mediator system. Six of the eight laccases demonstrated activity. One of these gut microbial laccases and three previously characterized laccases were then tested for their capacity to deplete 11 different food-associated chemicals. All four effectively depleted the agrochemicals cyflumetofen and fluazinam, as well as the industrial chemical bisphenol AF to a lesser extent. By linking global gut metagenomes with activity assays, this work demonstrates the untapped potential of mining human gut metagenomes for laccases and other microbial enzymes that can actively modify various agricultural and industrial chemicals.

**Importance:** As adverse effects of fluorinated compounds on human health are emerging, the responsible enzymes from the human gut microbiome of geographically diverse human cohorts mediating interactions with fluorinated compounds in the gastrointestinal tract remain poorly characterized. In a multi-study analysis of publicly available microbiome sequencing data, we linked the abundance, diversity, and phylogeny of laccases within the gut microbiome to the degree of urbanization of human cohorts, and experimentally demonstrated the ability of these laccases to deplete a range of food-associated fluorochemicals. Another significant contribution of our study is in the integration of rural catchment areas as a quantitative metric of urbanization in gut microbiome metagenomics and enzyme activity surveys.

## Introduction

Microbial biotransformation represents a critical yet insufficiently addressed factor in chemical risk assessment of fluorinated compounds, which have been linked with genotoxicity and endocrine disruption in humans (1–5). Fluorinated compounds are widely used in agriculture due to attractive properties such as increased stability, uptake and bioavailability (1, 6–9), and the share of fluorinated chemicals among newly-approved agrochemicals has risen from 9% to nearly 70% in the past quarter-century (3, 6, 10). While concerns have long been raised over the impacts of fluorinated pesticides on the environment and human health, transformation products can exhibit disparate toxicities compared to their parent compounds, potentially intensifying or mitigating environmental and human health risks (1, 5, 11–15). Gut bacteria have been shown to bioaccumulate and biotransform fluorinated compounds, with potential effects on bioavailability and side effects of fluorinated drugs (16–19). However, more research is needed on different classes of enzymes capable of fluorinated compound biotransformation, specifically their distribution, evolutionary relationships, and fluorinated substrate specificity. The human gut microbiome has been studied extensively in the past decade to understand its influence on human health, but the capacity of gut microbiota to act on fluorinated compounds and other environmental pollutants remains insufficiently explored (16, 20–22).

While the majority of gut microbiome samples sequenced so far originate from industrialized populations, studies have revealed that there are major differences in gut microbiome composition between industrialized and traditionalist societies (23–25). Lifestyle and urbanization play a crucial role in shaping the taxonomic structure (26) and functional gene composition of the gut microbiome, leading to differences in the resistome and metabolic capacity, e.g. carbohydrate active enzymes (CAZymes) (27). More recent studies have also explored the microbiota of intermediate cohorts which a large part of the world’s population belongs to, i.e., neither living in highly industrialized urban centers nor living traditionalist lifestyles such as hunter-gatherers (28, 29). Our study builds on these findings by investigating a class of CAZymes relevant for biotransformation of fluorinated compounds from the human gut microbiota along an urbanization gradient (30).

Specifically, our study focuses on laccases, a class of CAZymes previously shown to biotransform xenobiotics including fluorinated compounds (31). Laccases are multicopper oxidases (MCOs) encoded in plants, archaea, fungi and bacteria which are capable of oxidizing phenolic and other substrates with molecular oxygen as an electron acceptor (32, 33). The active site of MCOs, including laccases, consists of four copper (Cu) atoms of Type 1 (T1), Type 2 (T2) and Type 3 (T3) (34). Substrate oxidation occurs at the mononuclear T1 site, whereas reduction of oxygen occurs at the trinuclear site with two T3 Cu and one T2 Cu (35). Small-molecule redox mediators can also act as electron shuttles between the enzyme and the substrate. These mediator-assisted reactions significantly expand the oxidative capabilities of laccases, allowing them to target a broader range of substrates that are not directly accessible to the enzyme’s active site. Recently, we demonstrated the capacity of a bacterial laccase-mediator system to biotransform diverse xenobiotic organic compounds, highlighting their wide substrate range (33). Laccase homologs are widespread in environmental and host-associated microbiomes, yet their role in the human gut has not been defined, despite the intestine being a natural contact point between xenobiotics and microbial biotransformation enzymes.

We hypothesized that the diversity of laccase enzymes may vary in the gut microbiota of humans living in different locations and urbanization levels as a consequence of dietary and/or environmental exposures that shape the gut microbiome. Using publicly available metagenomic shotgun sequencing data from six different studies, we investigated associations of laccase-coding gene abundance and laccase sequence diversity and phylogeny with the degree of urbanization of sampling locations, classified according to urban-rural catchment areas (URCA) (30) as a quantitative measure of degree of urbanization. To our knowledge, this is the first microbiome study to use the URCA classification, allowing us to establish more objective and quantitative associations between microbiome and high-resolution geographical data studies across different studies and large geographic distances (30). We investigated the distribution and taxonomic lineages of laccase homologs in this multi-study analysis to differentiate putatively transient versus resident populations of the human gut. Using heterologous expression of diverse bacterial laccases, including eight derived from the human gut microbiome, we demonstrate that host-associated laccases depleted of several fluorinated compounds, suggesting that such reactions could also occur within the gut.

## Materials and Methods

### Study selection and metadata curation

Six studies (Table 1) on the healthy human gut microbiome with publicly available metagenomic sequencing data were selected for inclusion based on the availability of latitude/longitude metadata for the samples. The studies were intended to cover cohorts with a range of different lifestyles, including industrialized, subsistence farming and hunter-gatherer. Using the geographic metadata from the NCBI sequencing reads archive (SRA), each sample’s collection location was classified in terms of urbanization on a scale of 1 (most urbanized) to 30 (least urbanized) using the urban-rural catchment areas (URCA) approach (30, 36). Only metagenomes which could be successfully downloaded using q2-fondue were retained (37). In the Carter et al. (2023) study, in some cases multiple metagenomes originated from the same individual. For statistical tests where independent observations were necessary, the first collected sample from an individual was used.

**Table 1:** Studies with human gut microbiome metagenomic shotgun sequencing data used for our analysis, with sample countries of origin and total number of samples indicated for each study.

| **Paper** | **Country** | **# of samples (# of individuals)** |
| --- | --- | --- |
| Obregon-Tito et al. (2015) (25) | Peru | 17 (17) |
| Obregon-Tito et al. (2015) (25) | USA | 18 (18) |
| Rampelli et al. (2015) (27) | Italy | 10 (10) |
| Rampelli et al. (2015) (27) | Tanzania | 27 (27) |
| Lokmer et al. (2019) (38) | Cameroon | 57 (57) |
| Asnicar et al. (2021) (39) | UK | 997 (997) |
| Asnicar et al. (2021) (39) | USA | 97 (97) |
| Tamburini et al. (2023) (28) | South Africa | 169 (169) |
| Carter et al. (2023) (26) | Tanzania | 186 (74) |

### Bioinformatics

Metagenome data analysis was performed using MOSHPIT (40). Sequences were downloaded from the NCBI sequencing read archive (SRA) using q2-fondue (37). Raw sequences were trimmed to remove adapters and filtered with a phred score cutoff of 30 with fastp (41). Human-associated reads were removed by mapping to the GRCh38 reference human pangenome using bowtie2 (42). While this method may not completely exclude all human sequences, it was sufficient in the context of our study focused on laccase homologs (43). Reads were assembled into contigs with MEGAHIT using the meta-sensitive preset (44). Contigs were annotated functionally against the eggNOG database using DIAMOND (45, 46). Contigs with the eggNOG annotations “Cu-oxidase,Cu-oxidase_2,Cu-oxidase_3” for PFAMs and either “PFAM multicopper oxidase type” or “Multicopper oxidase” for description were considered to contain laccase homologs (47). Contig abundances were calculated according to the reads per million (RPM) method following read mapping using bowtie2 (42, 48). Contigs were classified taxonomically using Kraken2 (49). Strain isolation sources were retrieved for contigs which could be classified at the species level using BacDive and categorized as “Animal”, “Food”, “Human gut”, “Human other” or “Other” using a custom Gem created using the Google Gemini large language model (LLM) (50–52). Protein sequences coded by open reading frames (ORFs) were predicted from contigs using prodigal (53). Protein sequences were filtered such that only sequences with a length greater than 150 amino acids containing a start and stop codon as well as at least one copper binding motif (HXHG, HXH, HXXHXH or HCHXXXHXXXXM/L/F) (54). Protein Sequences were aligned using Clustal Omega (55). A phylogenetic tree of protein sequences was generated using IQ-TREE with ModelFinderPlus and bootstrapping with 100 iterations (56). Phylogeny aware distance metrics, namely Faith’s phylogenetic diversity and unweighted UniFrac, were calculated using the piccante R package (57–60). Associations between phylogeny and URCA classification were computed using Moran’s I and Blomberg’s K using the phylosignal R package (60–62). Contigs were binned into metagenome-assembled genomes (MAGs) with MetaBAT2 (63). MAG quality was assessed using BUSCO, medium- and high-quality MAGs with completeness >50% and contamination <10% were retained (64, 65). Taxonomy of contigs binned to medium- and high-quality MAGs was predicted using Kraken2, with MAGs being assigned the lowest taxonomic rank with over 50% of the contigs being assigned to it (49). A comprehensive overview of the bioinformatics pipeline for sequence processing can be found in Figure 1A.

**Figure 1:**
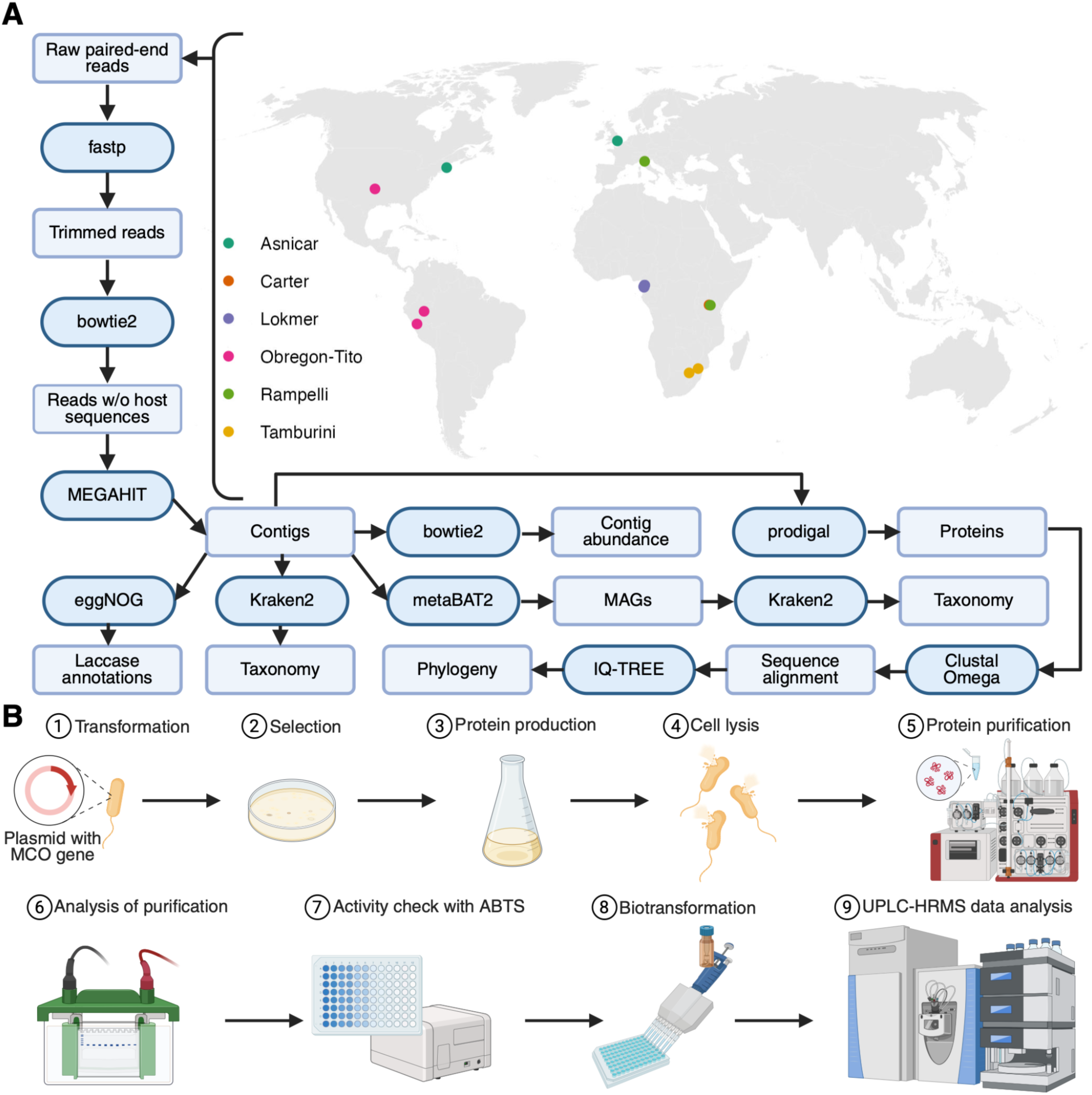
Overview of methods (**A**) Map of sample origin and overview of bioinformatics pipeline used for sample processing. Points on map are labeled according to study of origin as shown in Table 1 (**B**) Experimental methods used to assess enzyme activity

### Statistics and plotting

Statistics and plotting were done using R (version 4.5.1) within the R Studio integrated development environment (IDE) using the packages ape (version 5.8-1), ggnewscale (version 0.5.2), ggpmisc (version 0.6.3), ggpubr (version 0.6.1), ggtree (version 3.16.3), janitor (version 2.2.1), pacman (version 0.5.1), phylobase (0.8.12), phyloseq (version 1.52.0), RColorBrewer (version 1.1-3), readxl (version 1.4.5), rentrez (version 1.2.4), scales (version 1.4.0), tidyverse (version 2.0.0), treeio (version 1.32.0), ursa (version 3.11.5), and xml2 (version 1.5.0) (60, 66–84).

### Laccase selection and synthesis

All sequences and plasmids used in this study can be found in **Table S1**. Several ORFs representing laccase homologs from contigs assigned to the genera *Akkermansia* (Akk1 and Akk2), *Kurthia* (Kur), *Lactiplantibacillus* (Lac) and *Veillonella* (Vei1) were selected for synthesis as representative laccase homologs from our gut metagenomes. The full length Vei1 sequence was predicted by TMHMM-2.0 and Alphafold to contain an N-terminal disordered region (85, 86). Thus, a truncated version of the original *Veillonella* (Vei1_**Δ**120) was obtained. We searched for additional putative laccases associated with the gut microbiome using NCBI Blast (87). We identified a second *Veillonella* sequence with 66% aa sequence identity to Vei1 that also contained a predicted disordered region. Thus, a second truncated *Veillonella* (Vei2_**Δ**101) was procured. Two laccases encoded in *Enterococcus* species isolated from feces were selected (Ent1 and Ent2). Both source organisms are available from American Type Culture Collection (ATCC), ATCC 51266 and ATCC 6056 respectively. Finally, three additional non-gut reference bacterial and archaeal laccases (MCO1, aMCO, and mMCO) for comparison were selected, synthesized, and cloned as described in Yu et al. (33, 88).

### Laccase expression and purification

A full overview of experimental methods can be found in Figure 1B. All genes contained an N-terminal His-tag, and plasmids were transformed into *Escherichia coli* BL21(DE3) for heterologous expression. Single colonies were picked and inoculated into 5 mL of terrific broth containing antibiotic selection and cultured overnight at 37°C with shaking. For protein expression, a 400 µL aliquot of the overnight culture was inoculated into 400 mL of terrific broth supplemented with the appropriate antibiotic and incubated at 37°C with shaking at 200 rpm until the optical density at 600 nm (OD600) reached approximately 0.5-0.6. The protein expression was induced by addition of 1 mM isopropyl β-D-1-thiogalactopyranoside (IPTG) into the medium. To promote functional multicopper oxidase formation, CuCl_2_ was added simultaneously with IPTG at a final concentration of 0.25 mM. Cultures were then shifted to 18 °C and shaken for 4 h, followed by overnight incubation at 18 °C without shaking to establish microaerobic conditions conducive to active enzyme production.

After centrifugation, cell pellets were resuspended in 50 mL of lysis buffer containing 50 mM Tris-HCl (pH 7.5), 300 mM NaCl, 10–50% glycerol, 5 mM imidazole, and 1 mM PMSF (88, 89). Cell lysis was carried out using a French press (Avestin Emulsiflex C3) operated at 1200–1500 bar for approximately 5 minutes, or until the lysate became visibly less viscous and non-sticky. A sharp drop in pressure during operation typically indicated complete cell disruption. Vei1_Δ120 lysis buffer lacked PMSF and lysate was created using three separate one-minute passes through a sonicator at 50% power (Qsonica Q125).

Lysates were clarified by centrifugation before loading onto a 1.0mL Histrap FF column (Cytiva) using an ÄKTA FPLC system (Cytiva). The columns were washed (50 mM Tris-HCl (pH 7.5), 300 mM NaCl, 10–50% glycerol, 5 mM imidazole) and proteins were eluted using a linear imidazole gradient (20–500 mM imidazole in wash buffer). Eluted fractions exhibiting protein peaks were pooled and further processed by HiTrap desalting columns (Cytiva) with Sephadex G-25 resin to exchange into the final storage buffer (TNG buffer, 10 mM Tris-HCl, pH 7.5; 30 mM NaCl; 10% v/v glycerol) to remove residual imidazole. The purified laccases were aliquoted and stored at –80 °C until use. Concentrations of purified laccases were determined by Bradford assay (BioRad) or Pierce assay (Thermo Scientific).

### Colorimetric activity screening of selected laccases using ABTS assay in 96-well plates

Laccase activity was determined colorimetrically using the chemical mediator 2,2’-azino-bis(3-ethylbenzothiazoline-6-sulfonic acid) (ABTS) as a chromogenic substrate, following a previously described protocol (33). Briefly, enzyme activity was assessed in clear, flat-bottom 96-well plates. Standard 200uL reactions contained 100mM ammonium acetate buffer (pH 5.50), 500 μM ABTS solution, and 5 μL of purified laccase. The oxidation of ABTS to its radical cation was monitored spectrophotometrically at 420 nm using a microplate reader at 30 °C, with absorbance recorded every 2 minutes for up to 24 h. The molar extinction coefficient (ε) of ABTS radical at 420 nm was taken as 36,000 M^-1^ cm^-1^, and the optical path length for a 200 μL reaction volume was approximated as 0.56 cm.

### pH purification profile

The truncated *Veillonella* laccase (Vei1_Δ120) was used to generate a pH activity profile using the described ABTS assay. A stock buffer containing both 111 mM ammonium acetate and 111 mM MES was created to buffer between pH 4.0 and 7.0, and aliquots were adjusted with HCl to create solutions at 0.50 pH unit increments. 200 µL reactions were run in 96-well plates containing final concentrations of 4.0mM ABTS and 200 mM of the appropriate buffer. Assays were initiated with 1.2 mg of truncated *Veillonella* laccase and were monitored at 420nm for 10 minutes. Slopes for no laccase control reactions were subtracted from initial reaction slopes.

### Fluorinated chemicals selection

A set of fluorinated contaminants was selected (**Figure S1**) based on usage data in the fluorinated pesticide PFAS (Per- and polyfluoroalkyl substances) Data Hub and/or the USGS National Water-Quality Assessment Project (NAWQA) pesticide database, as well as their commercial availability and the availability of established detection methods by ultra-high-performance liquid chromatography coupled to high-resolution tandem mass spectrometry (UHPLC-HRMS/MS) (90, 91).

### High-throughput biotransformation experiments of fluorinated chemicals in 96-well plates

A mixture of selected fluorinated chemicals was used for enzymatic biotransformation screening. Reactions were performed in 96-well plates following our previously established workflow with slight modifications (33, 89). Briefly, individual fluorinated compounds were prepared as stock solutions (5–10 mM) in ethanol, depending on compound solubility, and stored at −20 °C until use. Prior to experiments, stock solutions were equilibrated to room temperature and sonicated for 10 min to ensure complete dissolution. Stocks were then diluted in ethanol and mixed to generate a working mixture containing each compound at 200 µM. For the screening experiments, 25 µL of the working mixture was first added to each well of a 96-well plate. The solvent was fully evaporated under room temperature. Subsequently, 125 µL of the buffer medium was added to each well, and the plate was mixed by pipetting up and down for approximately 1 min, followed by a 15 min incubation at room temperature to facilitate re-dissolution of compounds. An additional 100 µL of medium was then added, followed by a further 10 min incubation to facilitate complete re-dissolution of the compounds. Depending on the experimental condition, enzyme, mediator (ABTS), or nuclease-free water were subsequently added to reach a final reaction volume of 250 µL, corresponding to an intended concentration of ∼20 µM per compound. Prior to taking the first time point samples, wells were visually inspected for precipitation, and no visible precipitation was observed under these conditions. For the biological (enzyme-active) control, each well contained purified laccase enzyme, approximately 250 μM ABTS as mediator, and the fluorinated chemical mixture in 100 mM ammonium acetate buffer (∼pH 5.0). In the mediator-free enzyme control, the same components were included except for ABTS. For the abiotic buffer control, the fluorinated compounds and 250 μM ABTS were added without enzyme. Plates were incubated in dark incubator, shaking at 120 rpm, at 30 °C for up to 48 h, with samples collected at 0, 24, and 48 h to measure the parent compound removal by ultra-high performance liquid chromatography high-resolution mass spectrometry (UHPLC-HRMS/MS). Enzyme stocks used in this study differed in concentration and activity with ABTS (MCO1: 0.583 mg/mL; Vei1: 0.591 mg/mL; aMCO: 0.462 mg/mL; mMCO: 0.436 mg/mL). Approximately 3 µL of MCO1 and 20 µL of VEI, aMCO, and mMCO were added per well to achieve observable ABTS oxidation activity under the assay conditions. This approach was adopted because the primary objective of this study was to assess qualitative biotransformation potential across a broad set of compounds for different laccases, rather than to perform quantitative comparisons of enzymatic activity or normalize enzyme loading by concentration or activity. Accordingly, enzyme amounts were adjusted to ensure measurable reactivity within the experimental timeframe rather than standardized across enzymes.

### UHPLC-HRMS/MS analysis

The fluorinated chemicals were quantified using UHPLC-HRMS/MS (Q Exactive, Thermo Fisher Scientific). The analytical procedure was adapted from previously established protocols with minor adjustments, and full instrumental parameters have been described previously (88, 89). Briefly, for UHPLC analysis, 25 μL samples were loaded onto an ACQUITY Premier BEH C18 Column (particle size 1.7 μm, 100 × 2.1 mm, Waters) and eluted with nano-pure water (A) and methanol (B) (both amended with 0.1% formic acid) at a flow rate of 300 μL/min, with a gradient as follows: 95% A: 0 – 1.5 min, 95% – 5% A:1 – 7.5 min, 5% A: 7.5 – 9.5 min, and 95% A: .5 – 11.5 min. For HRMS, mass spectra were acquired in full scan mode at a resolution of 70,000 at m/z 200 and a scan range of m/z 100 − 1000 in positive/negative switching mode with electrospray ionization (ESI). Matrix-matched calibration was used to account for matrix-induced ion suppression or enhancement effects during electrospray ionization. Xcalibur 4.0 (Thermo Fisher Scientific) was used for data acquisition and analysis. Chemical structures were drawn, visualized, and annotated using ChemDraw Professional 20.0 and MarvinSketch (v19.20.0, ChemAxon, http://www.chemaxon.com). To assess compound removal, the incubation time at which each compound reached its maximum reduction across various control treatments was determined. Removal efficiency was calculated using Equation 1, where C_0_ represents the initial concentration and C_final_ denotes the concentration after incubation. Biological activity was considered significant when two criteria were met: (1) enzymatic treatment resulted in greater than 40% removal, and (2) removal in biological control was significantly (*p*<0.05) higher than that in the abiotic controls.

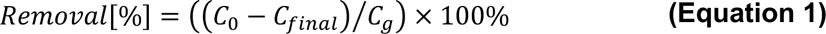

## Results

### Multi-study analysis and MAG recovery

To evaluate the global distribution of bacterial laccase homologs in the gut microbiome across diverse populations, we re-analyzed a set of six published studies from distinct geographic locations and lifestyles (Fig 1). These represent a gradient of urbanization following the URCA classification system, from urban centers (URCA 1-7) to peri-urban (URCA 8-13) to peri-rural (URCA 14-27) to hinterland (URCA 28-30), including hunter-gatherer communities. We recovered a total of 158,198 MAGs from our dataset, of which 48,312 were medium quality (completeness >= 50% and contamination <10%) and 17,626 were high quality (completeness >90% and contamination <5%) (**Figure S2**).

### Laccase-coding gene abundance as well as laccase protein sequence diversity and phylogeny correlate with urbanization

We examined the diversity of laccase homologs across human cohorts in our multi-study analysis, with the hypothesis that laccase homolog diversity and taxonomic affiliations will vary across lifestyle gradients. We found that the abundance of contigs containing putative laccase genes in the metagenomic samples correlates negatively with the URCA index for the samples (linear regression, adjusted R^2^ = 0.07, P < 0.001) (Figure 2A). This is a more pronounced effect size than for the housekeeping gene *rpoB* (linear regression, adjusted R^2^ = 0.02, P < 0.001) (**Figure S3**). Additionally, we used Faith’s phylogenetic diversity as an index of laccase genetic diversity (based on sequence similarity), demonstrating that the genetic diversity of laccase homologs in each sample with full laccase sequences is correlated negatively with the URCA index as well (linear regression, adjusted R^2^ = 0.12, P < 0.001) (Figure 2B). Unweighted UniFrac distance was calculated based on laccase genetic diversity and abundance in each sample to evaluate how laccase homolog composition varies across populations and URCA settings (Figure 2C), demonstrating a significant association with the URCA index (PERMANOVA, R^2^ = 0.07, p = 0.001). This further highlights that gut microbial laccases differ in protein sequence and abundance across urbanization levels.

**Figure 2:**
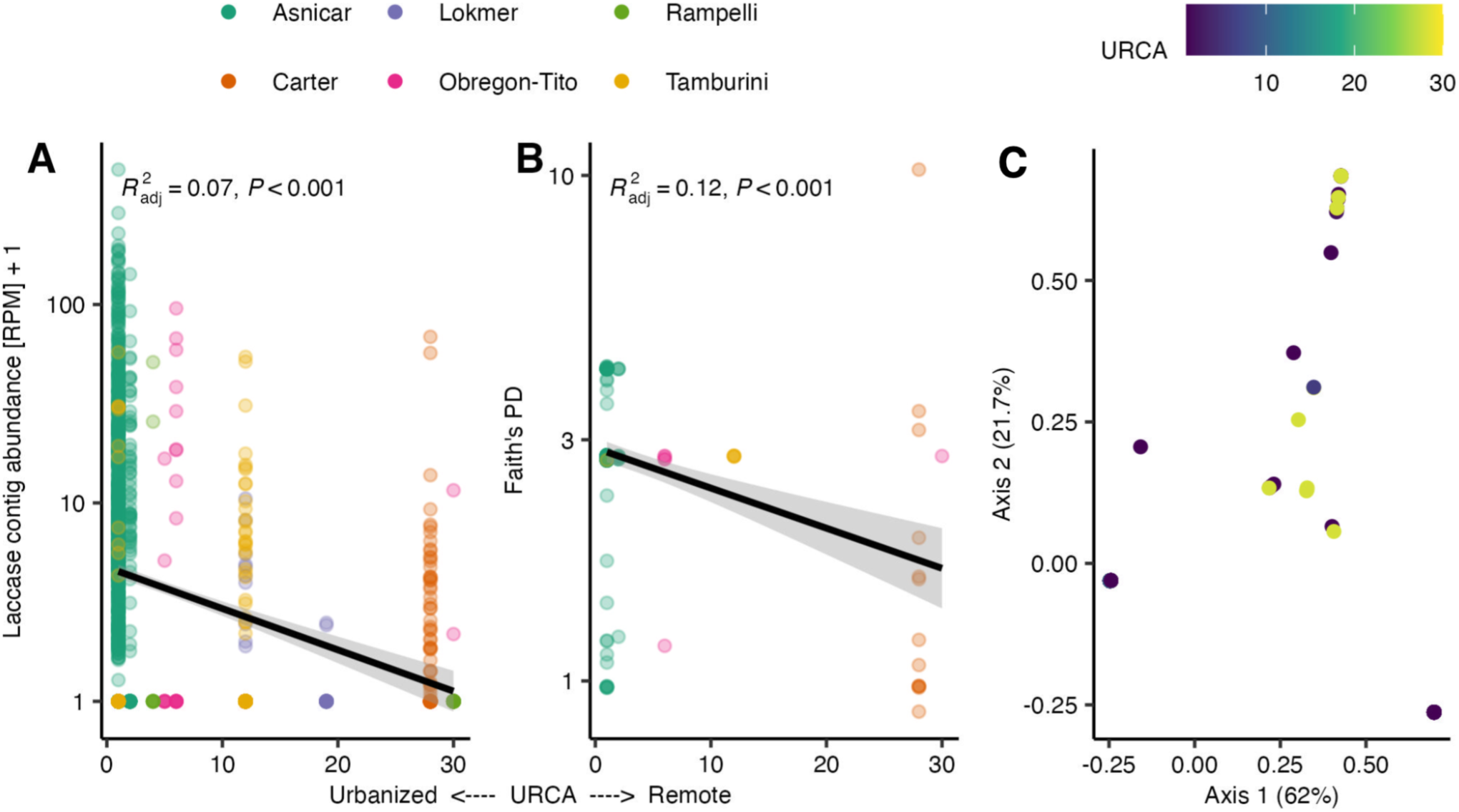
Laccase abundance and diversity metrics in association with URCA (**A**) Per-sample abundance measured as reads per million (RPM) of contigs containing putative laccase-coding genes plotted against URCA index values. A lower URCA index value indicates a greater degree of urbanization. Individual samples are annotated according to paper and country of origin. (**B**) Faith’s phylogenetic diversity of laccases in samples which contain laccase-coding genes plotted as a function of the URCA index (**C**) PCoA of Unifrac distances between samples based on abundance of contigs with laccase-coding genes and phylogeny of laccase protein sequences, indicating a significant association of laccase homolog composition and URCA index (PERMANOVA R^2^ = 0.07 p = 0.001).

As laccase homolog alpha and beta diversity correlates with URCA degree of urbanization, next we examined the laccase protein diversity in more detail to assess whether these differences were driven by differences in taxonomic composition. Construction of a phylogenetic tree of the laccases found indicates that the mean URCA of laccase origin sites correlates with phylogenetic distance (Moran’s I, I = 0.35, p = 0.001; Blomberg’s K, K = 0.0014, p = 0.003), indicating that laccases from cohorts with similar levels of urbanization share similar laccase clades and taxonomic origins as well. Many laccases found in samples from urbanized cohorts come from the genus *Akkermansia*, 47 from the species *Akkermansia muciniphila*. Overall, the laccases from (peri-)rural (URCA >= 14) samples come from more mixed taxa, with laccases found in 12 different families compared to 5 families in (peri-)urban samples (Figure 3) (30).

**Figure 3:**
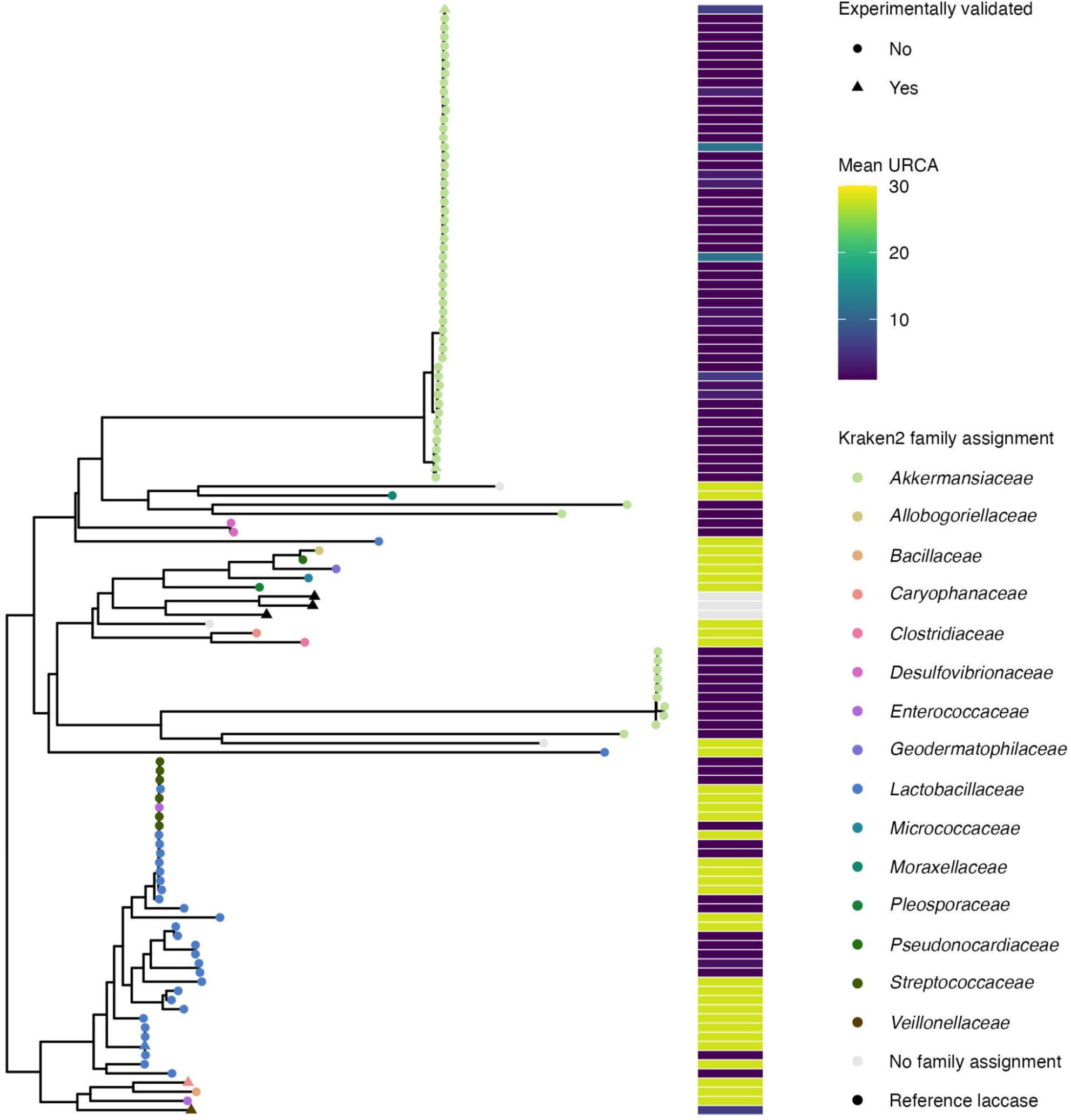
Phylogenetic tree of laccase amino acid sequences, with tips labeled according to family-level NCBI taxonomy classification assigned to contigs of origin using Kraken2, mean URCA index of all samples in which the laccase sequence was detected and experimental validation status.

Next, we examined the typical source location of bacterial species associated with laccase homologs in our multi-study analysis with the aim of distinguishing between putatively resident and transient members of the gut microbiota and identifying potential influences from food sources. Using the BacDive database, isolation sources of all strains associated with species with contigs coding for laccase genes were retrieved and classified as either animal, food, human gut, human non-gut or other (50). Notably, *Akkermansia muciniphila* is the only species with laccase coding genes where the majority of isolated strains originate from the human gut (**Figure S4**). This corresponds well with the fact that most medium- or high-quality MAGs that could be recovered and assigned a species-level taxonomy which contained laccase-coding genes were classified as *Akkermansia* at the genus level (**Figure S5**). A large number of bacteria detected in the gut metagenomes (6 out of 23 species containing laccase-coding genes) are primarily isolated from foods, namely *Furfurilactobacillus rossiae, Lactobacillus delbrueckii* subsp*. allosunkii, Lactococcus cremoris, Lentilactobacillus hilgardii, Limosilactobacillus pontis,* and *Weissella cibaria*, suggesting that these are more likely transient residents (**Figure S4**).

### Laccases from gut bacteria show activity with ABTS mediator system

As laccase activity has not been experimentally demonstrated from human gut bacteria previously, we selected several laccase homologs from our gut metagenome dataset to recombinantly express and purify for activity assays with an ABTS mediator system. Two laccase homologs from *Akkermansia* (Akk1 and Akk2) were selected as representatives of the *Akkermansia* laccase homologs found in the gut microbiome of urbanized cohorts, whereas a *Veillonella* laccase homolog (Vei1) was selected as an example found in gut bacteria of less urbanized cohorts. Additionally, we selected laccase homologs from the genera *Lactiplantibacillus* and *Kurthia* as examples from food- and animal-associated bacteria that are transiently present in the human gut as well as other environments. Two laccases encoded in *Enterococcus* species, isolated from feces, were also expressed, purified, and assayed.

The majority of His-tag purified gut laccases demonstrated activity with ABTS after one hour reactions compared to no enzyme controls (Figure 4A). Each triplicate reaction contained a fixed volume of purified enzyme (5 µL) rather than a fixed enzyme concentration because the purity of laccases varied as assessed by SDS-PAGE (**Figure S6**). A truncated *Veillonella* (Vei1_Δ120) showed the highest activity among gut laccases and SDS-PAGE revealed a clean band at the predicted 52.5 kDa size. The lowest activity was found in purified *Akkermansia* (Akk1), which encodes a membrane embedded C-terminal beta-barrel as predicted by Alphafold 3 (86). SDS-PAGE revealed little discernable purified Akk1 and a significant number of contaminating bands. We hypothesized that the beta-barrel could limit solubility and thus, activity might be present in the cellular lysate. Indeed, lysate from induced cultures expressing Akk1 showed significantly more activity with ABTS than uninduced samples, and SDS-PAGE shows clear induction of Akk1 (Figure 4B). A second *Akkermansia* (Akk2), which naturally lacks the C-terminal beta-barrel, and Ent2 did not show activity above baseline. Finally, we assessed the activity of Vei1_Δ120 at different pH values in an ammonium acetate and MES mixed buffer system. Vei1_Δ120 exhibited a pH optimum of 5 - 5.5 which is consistent with the mildly acidic environment of the duodenum and caecum (Figure 4C) (92).

**Figure 4:**
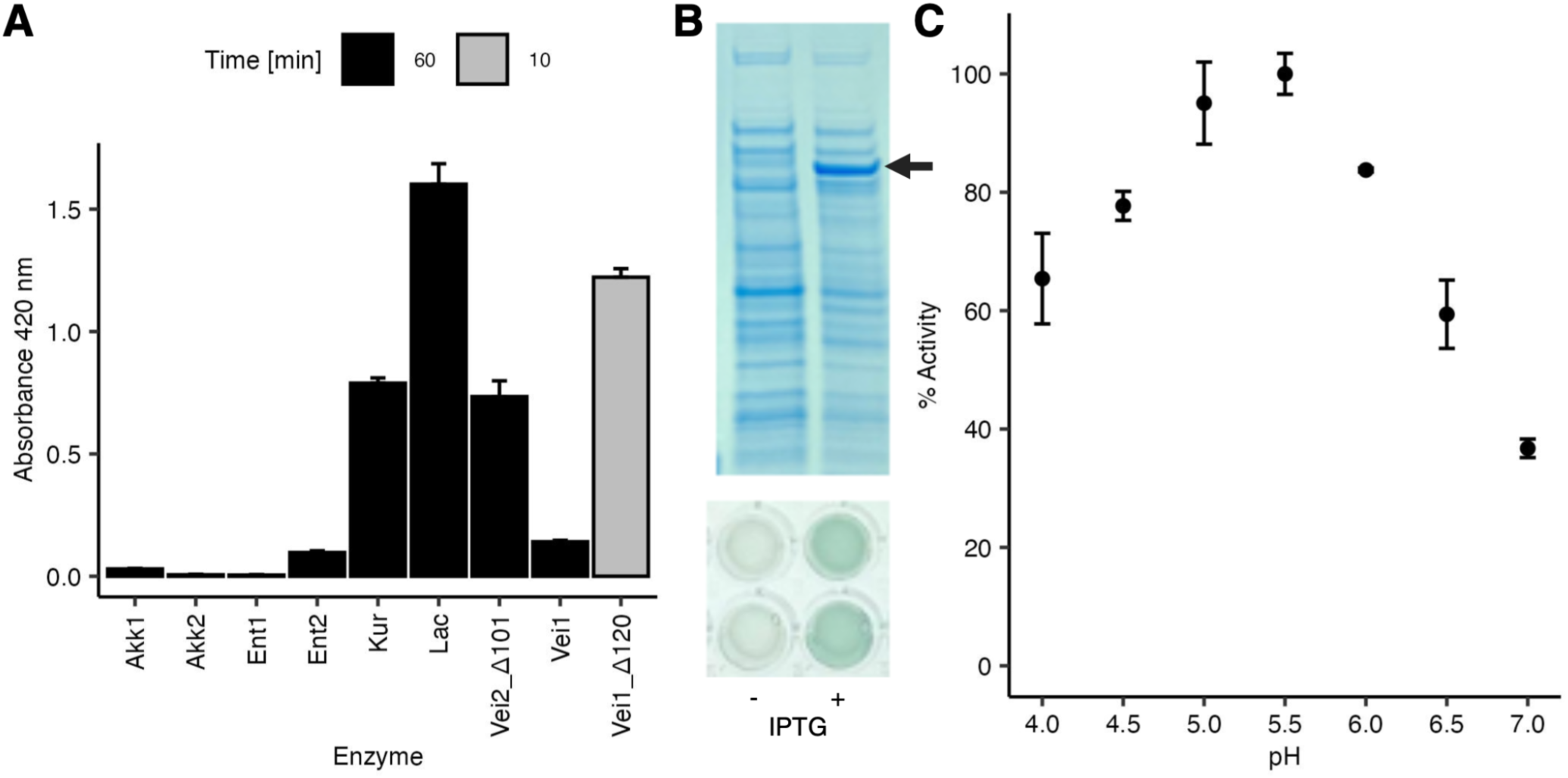
Verification of ABTS activity in predicted gut microbiome laccases. (**A**) Activity from 5 µL of purified laccases after 1 hr for all enzymes except Vei1_**Δ**120 (10 min). Activity for Ent1 and Akk1 is below 0.01 Abs/hr and is not considered significant. (**B**) Duplicate activity after 2 hrs from 5 µL of crude lysate of uninduced and induced Akk1 (indicated by arrow) expressed in E. coli. (**C**) pH activity profile of Vei1_**Δ**120 with ABTS.

### Depletion of selected fluorinated chemicals by representative laccases

To examine the activity of laccase-mediator systems with fluorinated chemicals, we evaluated the removal of a panel of 11 chemicals using 4 representative bacterial and archaeal laccases selected out of those synthesized to maximize diversity: Vei1 as a representative of our characterized metagenomic laccase homologs as well as an archaeal laccase (aMCO) and two bacterial laccases (MCO1, mMCO) (33, 88). As it has been well-characterized, MCO1 was considered as a positive control (33). Biotransformation experiments were conducted in a laccase-mediator-system using ABTS, a laccase-only system (without ABTS), and with abiotic controls to account for chemical losses in a laccase-mediator-system, pure enzymatic, and non-enzymatic losses.

In the laccase-mediator-system, two compounds, namely cyflumetofen and fluazinam, showed significant removal across all laccases compared to abiotic controls, indicating that laccases possess the capacity to mediate the transformation of these fluorinated compounds (Figure 5). Among the enzymes tested, MCO1 exhibited the broadest activity spectrum. In addition to cyflumetofen and fluazinam, this enzyme achieved greater than 40% removal of bisphenol AF. While the other three laccases also showed measurable removal of bisphenol AF, their activities were lower and did not consistently exceed abiotic controls to the same extent as the positive control enzyme. These results suggest that although laccases share a conserved oxidative capability, they differ in their capacity to transform specific fluorinated substrates. Such differences likely reflect variations in active-site structure or redox potential among enzymes from different ecological origins.

**Figure 5:**
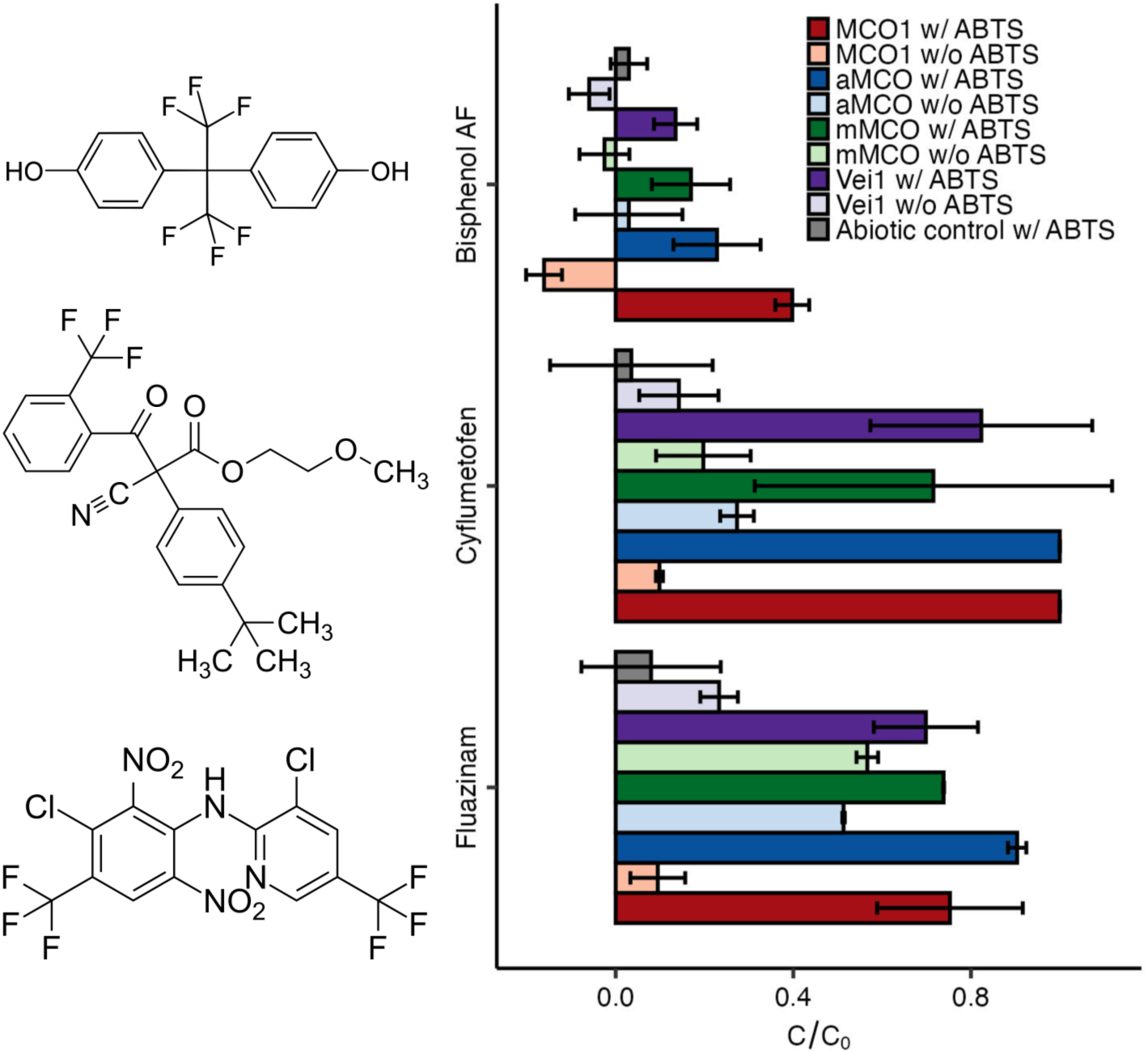
Removal of selected fluorinated chemicals by four representative laccase-mediator systems with the laccases MCO1, aMCO, mMCO1 and Vei1.

Furthermore, as ABTS is an artificial mediator and is unlikely to be present in natural systems, we assessed whether laccases could catalyze fluorinated compound transformation in the absence of an added mediator. As expected, overall transformation was reduced under mediator-free conditions (Figure 5). However, activity remained significant for specific enzyme–substrate combinations. Notably, both aMCO and mMCO showed measurable removal (>50%) of fluazinam without ABTS.

## Discussion

Fluorinated compounds have become widespread in food-contact applications in recent years, especially as agrochemicals and in food packaging materials (3, 93). In humans, the gut microbiome represents a major site for the biotransformation of xenobiotic compounds, including fluorinated agrochemicals (94–96). In this study, we characterized diverse archaeal and bacterial laccases including from the human gut microbiota of geographically diverse cohorts (31). We mapped the distribution of laccases in gut microbiota and experimentally validated their activity in mediator systems and role in fluorinated chemical removal.

Our results show that the gut microbiota of different human cohorts differs in terms of laccase abundance and diversity, implying that urbanization and lifestyle factors may influence fluorinated xenobiotic metabolism in the human gut. In line with studies showing that host lifestyles shape gut microbiome composition, we found that urbanization was associated with increased abundance of genes coding for laccases as well as increased Faith’s phylogenetic diversity (used here as a metric of laccase homolog sequence diversity, not community diversity) (97). In other words, both laccase abundance and genetic diversity varied across an urbanization gradient, suggesting that urbanization-associated factors may influence the spectrum of laccase activities within the human body, with wide-ranging effects as these encompass a range of enzymatic targets not limited to the fluorinated compounds tested in this study. A plausible explanation is that exposure to larger concentrations of xenobiotics (including increased dietary diversity) in urbanized populations leads to selection of a diverse array of bacteria capable of biotransforming these compounds (98–100, 96). Laccases can also oxidize plant structural components such as lignin, flavonoids, or tannins which are more abundant in non-industrialized diets (101–103). Thus, the trends observed in this study suggest that the effects of diverse dietary and/or xenobiotic exposures (and potentially other environmental exposures) in urban settings outweighs the effects of increased plant components on laccase abundance in human gut microbiota, even if it is not possible to link laccase abundance to exposure to one specific compound or group of compounds.

We also found a significant association between laccase phylogeny as well as UniFrac distances of laccases between samples with the URCA index. This suggests that the functionality (e.g., substrate range, performance kinetics) of gut-microbiota-derived laccases may differ between more and less urbanized cohorts, corresponding with established differences between other classes of enzymes in the gut microbiota of humans with different lifestyles, for example CAZymes (27). Although phylogenetic diversity was highest in urbanized populations, there was a notable diversity of laccases from less urbanized cohorts as a whole, including higher representation of unique laccase clades. Consistent with this finding, urbanized microbiomes are particularly enriched in laccase homologs originating from a single bacterial genus, *Akkermansia*. Many laccases were from the species *Akkermansia muciniphila*, which has been identified as being strongly associated with an industrialized microbiome (26). Conversely, the laccases from the gut microbiota of less industrialized cohorts originated from more varied taxa, including some associated with fermented foods, which aligns with previous studies indicating higher alpha diversity in the gut microbiota of rural cohorts (104).

Laccases from both human-gut-specialists (*Akkermansia*, *Enterococcus* and *Veillonella*) and animal- and food-associated genera (*Kurthia* and *Lactoplantibacillus*) demonstrated activity as part of laccase-mediator systems, representing the first report of laccase activity from these bacterial genera to the best of our knowledge. This is especially significant given that it represents the presence of oxygen-dependent enzymes in these mostly anoxic environments. (32, 105) Localized pockets of oxygen have been shown to exist within the human gut which would provide an environment where enzymes such as laccases could function (106, 107). Moreover, the apparent *K*_m_ for O_2_ for fungal laccases is in the range of 20-50 µM, indicating a high affinity for oxygen. Testing the affinity of laccases of gut microbial origin would further shed light on their potential to scavenge oxygen in oxygen-depleted environments (108). In addition to gut- and food-associated bacteria, we also describe activity of archaeal laccases, where previous studies have focused on fungal and plant laccases (101, 109). In short, our study significantly expands the range of organisms with characterized laccases, highlighting the potential for functional enzyme discovery from human and food microbiomes.

Regarding synthetic organofluorine compounds, we found that the gut *Veillonella* laccase, as well as three environmental laccases, were active in the removal of three fluorinated compounds, namely cyflumetofen, fluazinam and bisphenol AF, especially within a laccase-mediator system, whereas the remaining compounds were largely stable under the same conditions. These compounds are all aromatics with electron-withdrawing substituents including -CF_3_ groups, however this common backbone does not readily distinguish them from the other fluorinated agrochemical and industrial chemicals which were not transformed. Laccases preferentially oxidize substrates that can form stabilized radical intermediates, such as phenolic or aromatic amines, and are therefore more susceptible to single-electron transfer followed by coupling, rearrangement, or fragmentation reactions (109–113). In contrast, many fluorinated xenobiotics are dominated by electron-poor, highly substituted -CF_3_ motifs and lack readily oxidizable functional groups, which can make them poor laccase substrates even in laccase-mediator-systems (114–117).

Knowledge of biotransformations for these compounds is key for understanding potential adverse environmental or health effects associated with these compounds. Cyflumetofen, fluazinam, and bisphenol AF have been linked with various negative effects in humans and/or other animals, for example cytotoxicity or endocrine disruption (118–121). Gut biotransformations have previously been shown to alter the effects of fluorinated chemotherapeutic drugs through either deactivation or enhancement of cytotoxic effects indicate that the potential for microbial biotransformations to increase toxicity should be taken into account when evaluating the safety of food-associated fluorinated compounds as well (19, 122, 123).

In addition to our findings on laccases and their removal of fluorinated compounds, our study demonstrated a cross-cohort, multi-study approach to working with publicly available metagenomes by linking sequencing data to geographic metadata such as the URCA index (30). This approach allowed us to detect trends across cohorts from different studies and locations with high geographic resolution, whereas previous studies on urbanization and the gut microbiome have been limited to a single country classification systems to distinguish study populations (e.g., binary or trinary classification of urban and non-urban cohorts) (25, 27, 124). Future studies could expand on this approach to investigate other questions on environmental or host-associated microbiota, as well as integrating other relevant datasets such as chemical exposure data.

In conclusion, we demonstrate that phylogenetically disparate laccases are widespread in diverse human cohorts and capable of depleting various food-associated fluorinated compounds. This lends deeper insights into how human lifestyle factors relate to the prevalence of pathways for metabolism of xenobiotic fluorinated compounds. Together, these findings expand knowledge on the substrate range of laccases to include fluorinated compounds of public health relevance, link patterns of laccase abundance and diversity to human geographical factors and highlight potential approaches for further studies with publicly available microbiome sequencing data. These outcomes point to the possibility for microbial biotransformation of fluorinated food contact chemicals, with potential health risks that should be taken seriously in chemical safety assessments. Further research to assess *in vivo* activity and consequences is warranted, as is further investigation of a larger range of enzyme classes capable of fluorinated compound biotransformation.

## Data Availability

Sequencing data are reused from NCBI SRA (https://www.ncbi.nlm.nih.gov/sra) and are associated with the following BioBroject accessions: PRJNA268964, PRJNA278393, PRJEB27005, PRJEB39223, PRJNA678454, PRJEB49206 (25–28, 38, 39). Code used to generate figures is available under https://github.com/MSM-group/gut-laccases.

## Acknowledgments

N.A.B. and S.L.R. acknowledge funding from ETH Research Grants (23-2 ETH-047) and the Uniscientia Stiftung. Y.Y. acknowledges support from the Eawag Academic Transition Grant and Stanford Plant-Based Diet Initiative (PBDI), with the content solely the responsibility of the authors and not necessarily reflecting the official views of the PBDI. S.L.R. acknowledges support from the Swiss National Science Foundation (PZPGP2_209124). The work (proposal: 10.46936/10.25585/60008420) conducted by the U.S. Department of Energy Joint Genome Institute (https://ror.org/04xm1d337), a DOE Office of Science User Facility, is supported by the Office of Science of the U.S. Department of Energy operated under Contract No. DE-AC02-05CH11231.

