## Supplementary Information for "Laccase-mediated biotransformation potential for fluorinated compounds by geographically diverse human gut microbiota"

**Table S1**: Overview of laccases tested in this study.

| Name | Sequence | Vector | Antibiotic | Cut sites | Scientific name |
|---|---|---|---|---|---|
| Vei1 | MKKYSNYMKALLVAVAITGGT<br>AIVSAHDTVTTNDCCYDSATA<br>TNSQNMTANMSSMMGGMVN<br>NMMNNQSSANNSSMHSMMN<br>GQNSMDSNSMHSMMSGMMN<br>GGMMGMTATNSMNGMMGM<br>MGHMHGVDDVPALKQVQQAA<br>KPLAIPPMQKGTVDENGVRHL<br>EVTAQEGKTAIKDGALTDTYG<br>YNGSVLGTTLYMKHGEKVAIK<br>LKNELPEKTTYHWHGMVVRS<br>DVDGGPHYPIAANGGTGEVEF<br>TVDQSANTAWYHPHAMGLTA<br>SQVYKGLAGFIYVDDNKEGAL<br>NLPHTYGVDDIPVAIQERNFTA<br>DNQWDYEKDYNADGVYGDTL<br>VVNGTINPYFDVKTNMVRLRL<br>LNGSNARTYTLQLSDMSPMM<br>QVAGDGGILPWPVPKQSITIAP<br>GERVEVLVDFSKYKDALPSLV<br>TTNGTNGTANVLNFKRGADTL<br>TAEAKVNTDMAAWNYNNMNA<br>SDDLTKPADKSIVMSGMAQNV<br>MINGKKFDMGRIDFTSKLGST<br>EIWDVSNADPGMMGMMHPFH<br>VHGVQFEVLSRNGRPVDAAE<br>KGLKDTIQVDPGEHVRIRLHFT<br>KPGVFMVHCHILEHEENGMML<br>QLEVK | pRSETA (full sequence)<br><br>pET28a+ (Δ120) | Amp (full sequence)<br><br>Kan (Δ120) | EcoRI/HindIII (full sequence)<br><br>Nde1/Xho1 (Δ120) | *Veillonella* |
| Vei2 | MKINQMKLAKLALAVTLMTGS<br>GYVVNAQMMNGMMNQSMMS<br>QDVMDQDSMNSMMNTMMGQ<br>SSSNQNYGNGMMNGSMMNG<br>MMNQNSMHNTMMNNMMSG<br>QGMMGMHDEAMTLHQSTQA<br>PRPLAIPPLVKGTPDGNGRMV<br>YNVTAQTGESDIKDGQKTATY<br>GYNGNVLGPALLMKRGEKVTI<br>NLTNNLPEATTFHWHGLVVRS<br>DVDGGPHYPIAANGGTGKIEF<br>TVNQPAGTAWFHPHAMGSTA<br>SQVYKGLAGLIVIDDNEIGKLN<br>LPHEYGVDDIPVAIQDRLFTKD<br>NQWDYKSAYSDDGVYGDTLV<br>VNGTINPYFEVPKKLVRVRLLD<br>GSNARNMTLRLSDGSTMYQIA<br>SDGGLLEAPVATQEITLVPGE<br>RAEVLIDFSRYANADKLPSLVT<br>QNGTINVLEFRKAASWNTDSA<br>ASVNNTIPTWNFETKKHDEEL<br>QKAATTDKIAQHIVMSGMAKD<br>VTINGKKFSMDRIDLTAKLGST<br>EVWEISNSDGMMGMIHPFHIH<br>GVQFEVVSRNGVPTTPDERG<br>LKDTISVAPGENVRIRIHFTQP<br>GIFMVHCHILEHEENGMMLQL<br>QVK | pCDF-Duet-1 (Δ101) | Spec (Δ101) | BsaI/BsmBI (Δ101) | *Veillonella* (Δ101) |
| Ent1 | MKNIRTSFSDSPSKETMQEER<br>PNRNTMGNMMTNEQNIELKNI<br>SQSEQPLLIPPLLEPGSEDDEN<br>ISYNITAKEGEFQIKEGKKTKTL<br>GYNRDFLGPVIRMKQGQKVTL<br>TTTNQLKSKTSFHWHGLKIPS<br>EVDGGPHQVVEPGEKKVISFT<br>VDQDASTLWFHPHPEGETAS<br>QVYRGLAGLMYIDDENSDSLN<br>LPNNYGKDDIPLIVQDKSFDSD<br>NQIDYETDFYSDGTQGDTLLV<br>NGTINPYIEVNSRWMRYRIISG<br>SNSRNFTFSLKDNQPFYQIAS<br>DGGLLNSSTKLTSLTLSPGER<br>AEILVDTNDYKNGDTLKLLANN<br>TVALTMKINKNKSNTTFEPSKK<br>LNSIKDIDTENLDKLDRQKIDLK<br>GMSHMVSINNKQFDADRIDLY<br>KRVNTQEVWEVNNISGMMGG<br>MIHPFHIHGVQFQIISRNGQLP<br>QANERGWKDTVLVNPEETVE<br>LLVEFDHEGVFMYHCHILEHE<br>EYGMMGQMEIK | pCDF-Duet-1 | Spec | BsaI/BsmBI | *Enterococcus dispar* ATCC 51266 |
| Ent2 | MQQNKEEHAEVVFQEEQRQI<br>SSRMSNMMGDNQEADLKVVH<br>QTERQLLIPPVLEPSSKSKENV<br>IYDIVAQNGEVQIMDGEKTETL<br>GYNGDFLGPVIRLKKGQKVTIN<br>TTNNLDASTSFHWHGLKVASD<br>ADGGPHQIIEAGQKKSVTFEV<br>DQEASTLWFHPHSEGETASQ<br>VYKGLAGLMYIDDDNSKSLDL<br>PSKYGVDDIPLIVQDKSFSSTN<br>QINYENDFNSDGTKGETLLTN | pCDF-Duet-1 | Spec | BsaI/BsmBI | *Enterococcus durans* ATCC 6056 |

| | | | | | |
|---|---|---|---|---|---|
| | GTINPYIEIKNRWMRYRIVNGS<br>NSRNFTFNLDNDESFYQIATD<br>GGFLNTSVKLSKLLLAPGERA<br>EILVDTQNYKKGKVIHLLANNL<br>VALTMRIENTIDNKEFNPSDSL<br>NTISTLDEKKLEDLTRQSINLS<br>GMSHMVNINNKQFDMERIDLY<br>KKLGTQEIWEVNNISSMMGG<br>MIHPFHIHGVQFQILSRDGNQP<br>ALNEQGWKDTVLVNPDEIVEL<br>LVKFDREGIFMYHCHILEHEEY<br>GMMGQMEIK | | | | |
| Kur | MKKTLIASLLVGTALLAACGNE<br>DGATDHTTGASEHQHSMANT<br>HQNGSMDHEQLVLKDNKGTN<br>TLIFPAILKPDSEKGKNVEYSLT<br>AQAGTTELFNGYETKTYGYNG<br>NLLGPTLRLKEGQHVTIHLKND<br>LPEATTFHWHGLEVSGKADG<br>GPHALIEPGEEKTIQFTVTQQA<br>ATLWYHPHPMGNTAQQVYKG<br>LAGLLYIEDDNVEALNLPNDYG<br>KNDFPIILQDRAFVEGKQLDYS<br>KVANSDGTYGDTMMINGVINP<br>VLKTDKKLVRLRILNGSNARNY<br>NVHFDNDMAFTQIATDGGFLN<br>KPQKMKELLLGAGERAEILVDL<br>SKVKGKRVSLVNDQNVVLLPI<br>DVSDTKSAVNASTKLNTIKVDK<br>ALLEQQPTKVVKMEGMRENV<br>TINGKKFDEKRVDFTQKKGET<br>EVWEIDNAITDEGGMIHPFHIH<br>GTQFLVLSVNGEKPEPSLQGY<br>KDTITLQPGQKARIAVKFPYEG<br>MYMFHCHILEHEDNGMMGQI<br>EVK* | pCDF-Duet-1 | Spec | BamHI/KpnI | *Kurthia* |
| Lac | MAKKVYTDYFFDEPAYNTHDG<br>GYIPLVTPKVEPQPLAIPPLLKP<br>DRQTDTDDYYTVTAQESETQF<br>LPGKKTKTWGYNAGFLGQTIV<br>FRNGKQTHIDLENKLPELTTFH<br>WHGLNVPGPITDGGCHAPVY<br>PGETNHIDFKVHQPAATTWLH<br>AHPCPSTATQVWKGLATMVII<br>KDDVEDQLPLPRNYGVDDIPL<br>VLQDREFHDDNQFDYRADYD<br>PDGVQGHTALVNGTVNPYFD<br>VTTQRVRLRILDGSNRREWRL<br>HFNDDLEFAQVASDGGILPAP<br>VYMTKVMMTCAERDEIVVDFG<br>QYQPGDEVTLMTDDTPLCRF<br>RIKSFVPDDTKLPEHLVDIPDE<br>TPTPDLPVRTITMDGMDDEVA<br>LDGKKFDMSRIDARQKVGDVA<br>IWEIRNTNSTENGMVHPFHVH<br>GTQFRVLARNDGPVYPNEHG<br>LKDTVGVNPGETVRIKVKFELT<br>GVYMYHCHIIEHEDGGMMAQI<br>ESYDPQHPQTYHLMDMDTLR<br>NAFAKEQGIKPEDVWMPGM | pCDF-Duet-1 | Spec | BamHI/KpnI | *Lactiplantibacillus* |
| Akk1 | METVEYDLYVRNTPVNFTGTA<br>RPAMSINGSIPGPVLHFTEGDT<br>AVIRVHNMMDTETSFHWHGLL<br>VPNDQDGVPYLTSAPVKPHTT<br>HTYTFPIIQNGTYWYHSHSGL<br>QEQSGLYGAFVVHKRRNDPA<br>RRAEDTLPEYTLVLSDWTNEN<br>PNEVNRKLHTGSDWFSIRKGS<br>VQSYWEALRAGYVGTKLTSE<br>WKRMNPMDVSDVYYERFLLN<br>GSPHASLPRLKAGDRVRLRIV<br>NGASSTYFWLRYSGGKIRVVA<br>SDGKDVVPVDVDRMIIAVSET<br>YDVILTVPETGRAFEFQATAED<br>TTGSSSLWLGHGEKQPLRPFP<br>KLNYFKNMKRMNGMMTMGG<br>SMKMMTMSSGSMPGMKHMN<br>HNMSGGMDSPRAGGMHMMS<br>MSSGSSHGKHHAEDMQEDE<br>GEVTLTYDMLRSLAKTNLPSG<br>VPVKELHFQLSGNMNRYVWSI<br>NGRTLSETDRIMIREGQNVRIIL<br>TNNTMMRHPMHLHGHFFRLV<br>NKQGDFSPLKFTVDIQPMATQ<br>VIEFNAAEKTRGNWFFHCHILY<br>HMMSGMGRIFTYEDSPPNPQ<br>LPHPRQALQHVYAMDRKWYL<br>TVNNDFASNGNIGDLEFGGTR<br>WSIQGEWQTGYKDTRGYEAE<br>ARLGRYIGEKQWLYPYIGMDW<br>TYRKGEAGERNMFRQTTRKD<br>RELDGTLGTRYTLPLLLVADAR<br>IDTDGKVRLQLERDDIPLTSRL<br>RLSFSLNTDRDYSVGLHYILTP<br>HVSISTNYDNNLHWGVGLMLT<br>Y | pCDF-Duet-1 | Spec | BamHI/KpnI | *Akkermansia* |
| Akk2 | MKTVEYDLYVRNAPVNFTGVT<br>RQAMTINGNIPGPTLHFTEGD<br>TAVIRVHNLMDTQTSFHWHGL<br>LVPNDQDGVPYLTSAPVKPHT<br>THTYTFPIIQNGTYWYHSHSG<br>FQEQSGLYGAFIVRKRPDDPA<br>RRTEDALPEYTLVLSDWTNEN<br>PQEVNRMLHTGSDWFSIRKG | pCIOX | Kan | BamHI/HindIII | *Akkermansia* |

| | | | | | |
|---|---|---|---|---|---|
| | SVQSYWEALRSGCMATKLLS<br>EWKRMNAMDVSDVYYECFLL<br>NGIPRASLPRLRGGDRIRLRIV<br>NGASSTYFWLRYAGGKIRVVA<br>SDGKDVVPVDVDRMIIAVSET<br>YDVILEVPESGKAFEFQATAED<br>TSGSSSLWLGHGERQPLRPF<br>PKLNYFKKMKQMNGMMTMG<br>GNMKMHHHGMPTAHSGNMGI<br>MDMKSGSSHGGGHGNMQED<br>GEGTTLTYDMLKSPSRTTLPS<br>GVPVKELHFMLTGNMNRYVW<br>SINGKTLSETDRIMIREGQNVR<br>IILTNNTMMRHPMHLHGHFFR<br>LVNRHGDFSPLKFTADIQPMD<br>T | | | | |
| MCO1 | MKKTDNDTTQDSTPDSPSNE<br>SRRAFLKTTGATVIAAPAILTSR<br>KSMAQVVPGVAPSPAT<br>PWQVDLPDAIEPLQPTDLFTD<br>AYRIPQGTANTAYGECGRNRH<br>QRWNDFFGNPQLQPGQADTY<br>ELRVKEDNNYSFHPDYPDQLA<br>WTYEGVDGNGQPYHNPTFFA<br>RYGRPVIVRLHNELPKDHEGF<br>GTPEISMHLHNLHTPSESDGF<br>PGDYFSPYKAGPTLTKPGNFK<br>DQFYPNIYAGLDEFPKDVDNP<br>VGGDKREALGTLWYHDHTLD<br>FTAANASRGMAGFYLLFDDLD<br>SGDENDPNPAALRLPSYPYDY<br>PLLFQDKRFDASGIHIYDQIDP<br>EGTLGDKITVNGKIEPVLKVAK<br>RKYRLRLLNAGPSRYYEFYLV<br>NAAGTPQTFDYIANDGNLLPT<br>TLRNQTRVRLGVAERGDIVVN<br>FNRYALGTELYLVNRLVQTST<br>RGPGAVQAPGDRVLKIVVNRT<br>AADNSQVPNTLRPIRRPTAEEI<br>ASAPVRRWTFARTSGLWTINS<br>KLVNVKSPAARISKESMEIWDI<br>SNPNGGWSHPVHIHFEEGHIL<br>QRFVNGAAVAVPAHERGRKD<br>VYNLGPNERIRLFIRFRDFPGK<br>YVMHCHNMIHEDHAMMVRW<br>DIVE | pRSET A | Amp | EcoRI/HindIII | *Nitrosomonas* |
| mMCO | MTTAAAAGAPLILTSRKSGAQ<br>VVVPPSPPTTPWVQELPRAIT<br>PLAAVPSLSPAPTLTANTLGGE<br>CGRAEHQRFAELTAPGVCGP<br>PELYEMFARENPGWVFNPAY<br>PPQPVWGFSGSAADPATSPG<br>PTLFGHYGHSVICRIHNQLPAN<br>HVGFGTPEISTHLHNLHTPSES<br>DGFPGDFYSATIAGPTLGAPG<br>QFQDHFYPMVYAGLDTYGGIG<br>DSREALGTLWYHDHTMDFTA<br>PNLVRGMAGFYLLFDDLDTGN<br>EQTGLRLPSHPYDYPIFLQDK<br>RFDANGILTYDQFDPEGTLGD<br>KVTVNGKIEPVLRVERRKYRF<br>RFLNGGPSRFYEVYLQNRGAT<br>SVYTYTYIANDGNLLPAPLLNQ<br>FRFRLGVAERAEIVVDFARFPL<br>GTELYFVNQLLQDNTRGPGNV<br>RAPGTRLLKIIVDRNPAQPDTS<br>RVPSALRPLPPIPTDFSGIPVR<br>RWVFERSGGMWSVNGQFFN<br>VNVPRATIAKGAGEIWELVNP<br>ENGWEHPVHIHFEEGRIIEKRV<br>NGAVVPIPLHERGRKDVYTLG<br>KDVSFKVFLRFRDFKGKYVMH<br>CHNLIHEDHAMMVRWDIV | pRSET A | Amp | EcoRI/HindIII | *Methylmirabilota* |
| aMCO | MKKTDNDTTQDSTPDSPSNE<br>SRRAFLKTTGATVIAAPAILTSR<br>KSMAQVVPGVAPSPATTPWQ<br>VDLPDAIEPLQPTDLFTDAYRI<br>PQGTANTAYGECGRNRHQR<br>WNDFFGNPQLQPGQADTYEL<br>RVKEDNNYSFHPDYPDQLAW<br>TYEGVDGNGQPYHNPTFFAR<br>YGRPVIVRLHNELPKDHEGFG<br>TPEISMHLHNLHTPSESDGFP<br>GDYFSPYKAGPTLTKPGNFKD<br>QFYPNIYAGLDEFPKDVDNPV<br>GGDKREALGTLWYHDHTLDF<br>TAANASRGMAGFYLLFDDLDS<br>GDENDPNPAALRLPSYPYDYP<br>LLFQDKRFDASGIHIYDQIDPE<br>GTLGDKITVNGKIEPVLKVAKR<br>KYRLRLLNAGPSRYYEFYLVN<br>AAGTPQTFDYIANDGNLLPTTL<br>RNQTRVRLGVAERGDIVVNFN<br>RYALGTELYLVNRLVQTSTRG<br>PGAVQAPGDRVLKIVVNRTAA<br>DNSQVPNTLRPIRRPTAEEIAS<br>APVRRWTFARTSGLWTINSKL<br>VNVKSPAARISKESMEIWDISN<br>PNGGWSHPVHIHFEEGHILQR<br>FVNGAAVAVPAHERGRKDVY<br>NLGPNERIRLFIRFRDFPGKYV<br>MHCHNMIHEDHAMMVRWDIV<br>E | pRSET A | Amp | EcoRI/HindIII | *Thermoproteota* |

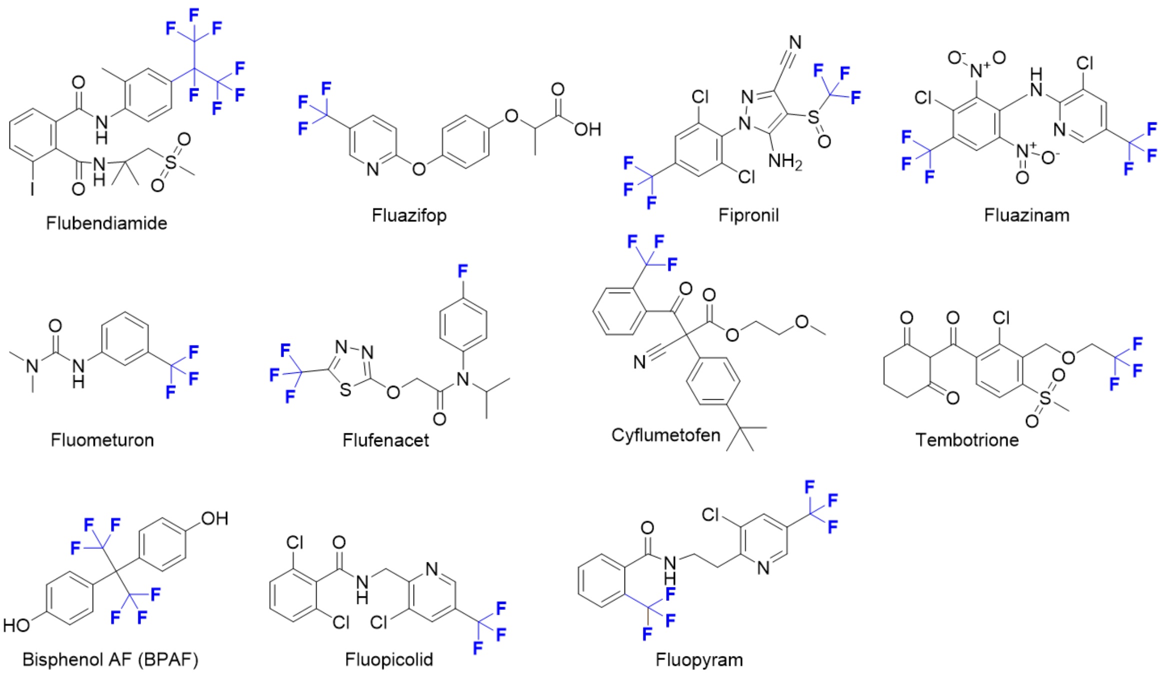


**Figure S1**: Chemical structures of the 11 selected fluorinated compounds used in this study.

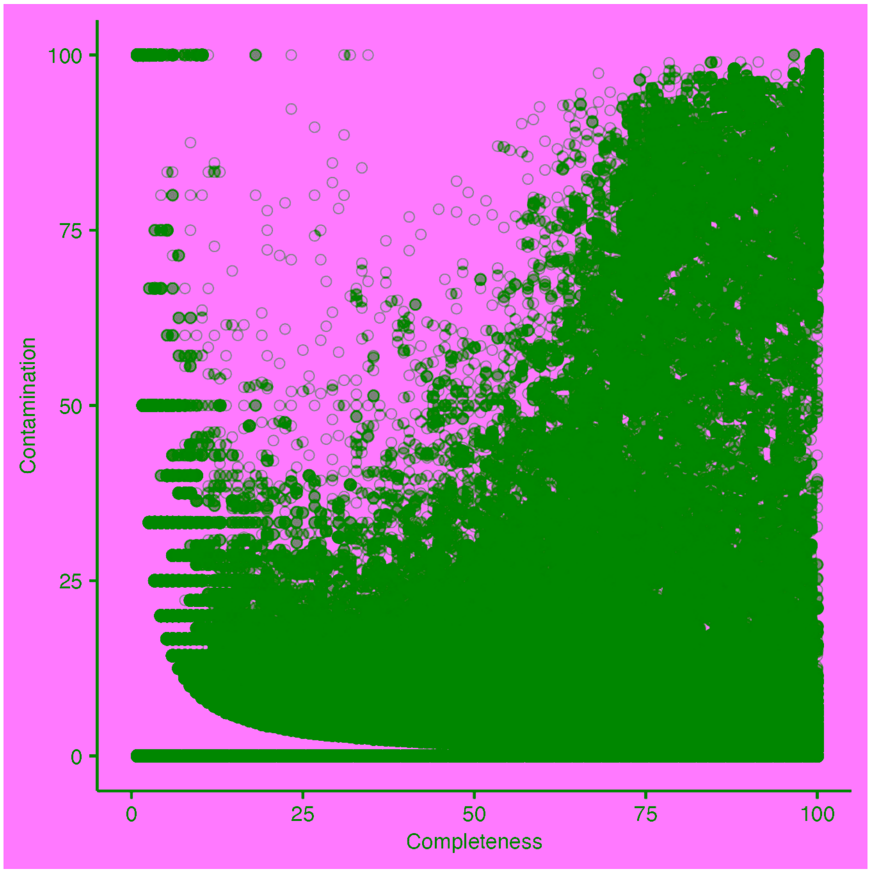


**Figure S2:** Completeness and contamination of recovered MAGs as evaluated by BUSCO

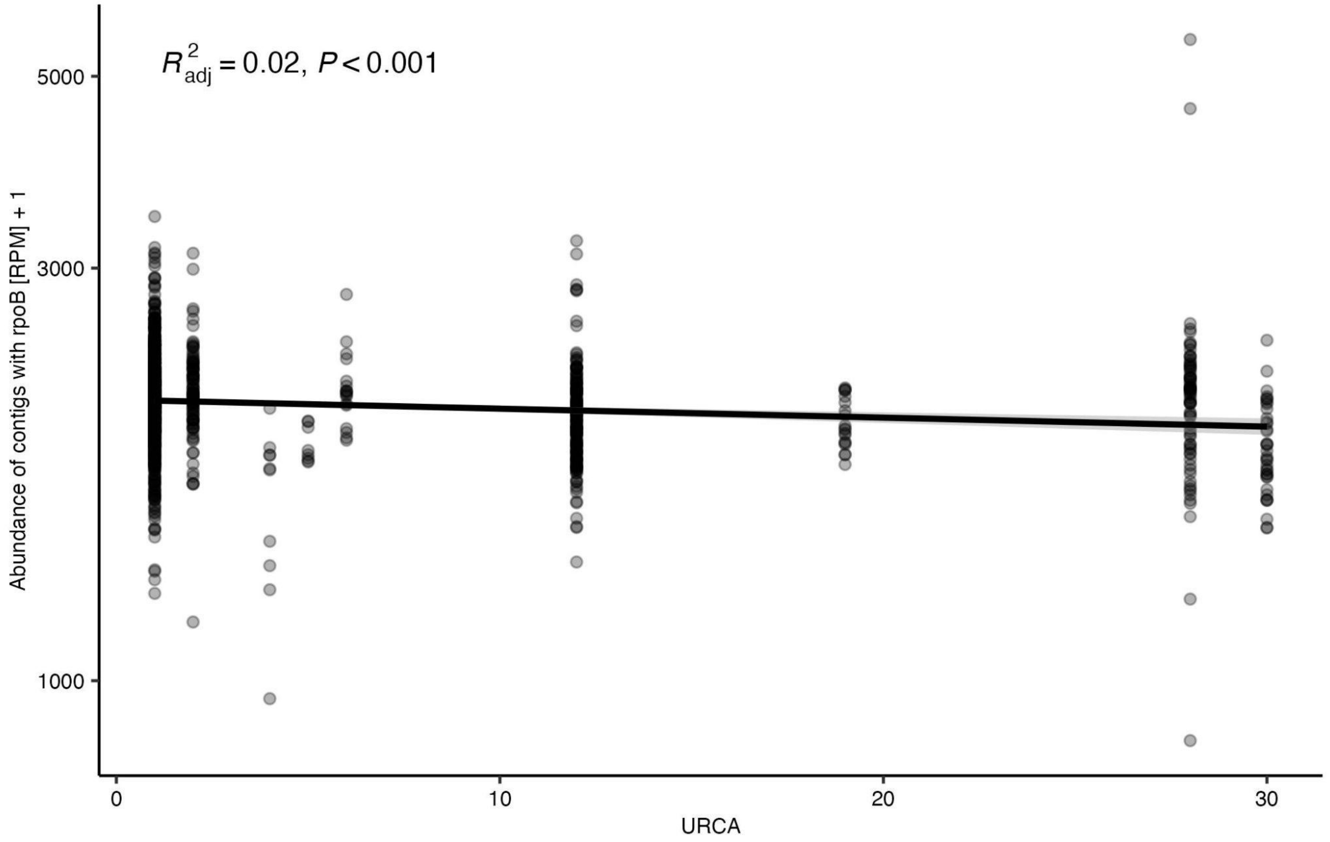


**Figure S3:** Per-sample abundance measured as reads per million (RPM) of contigs containing putative *rpoB* genes plotted against URCA index values. A lower URCA index value indicates a greater degree of urbanization.

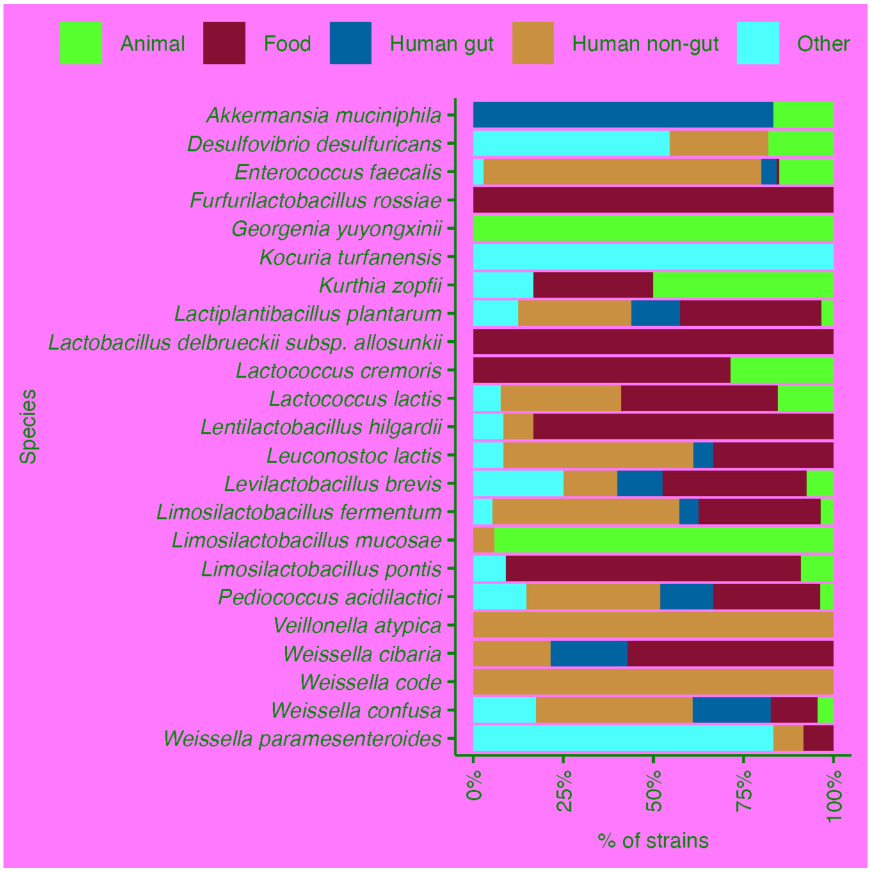


**Figure S4:** Percentage of strains within the BacDive database isolated from different sources for species where laccase-coding contigs were found. Strain isolation sources were categorized as animal, food, human gut, human non-gut or other.

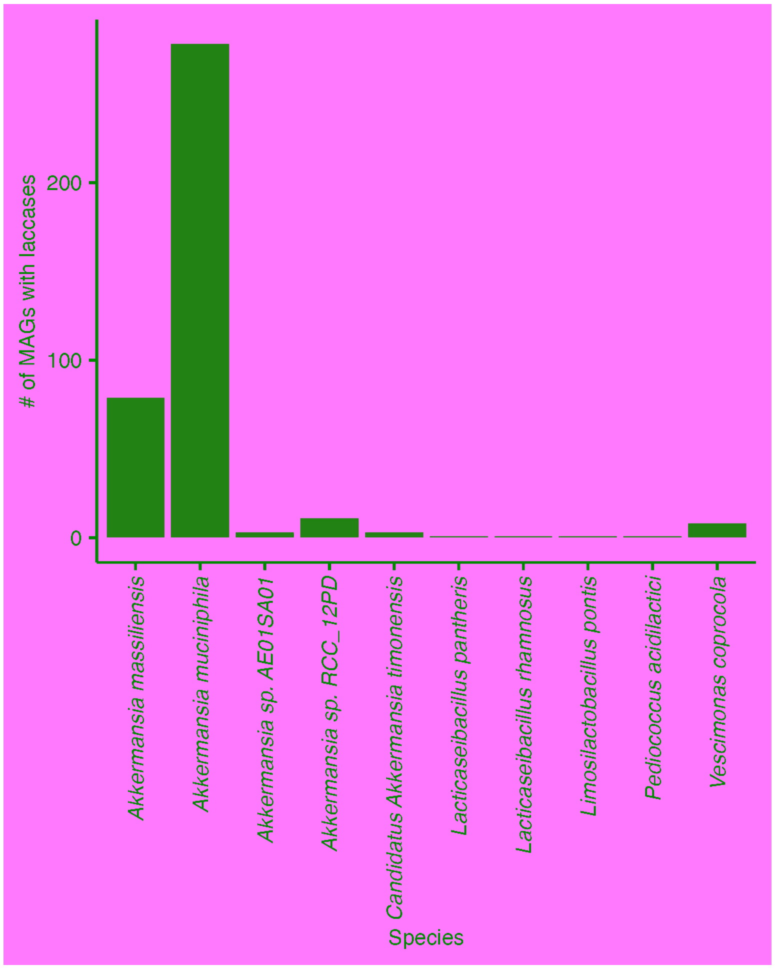


**Figure S5:** Taxonomic classification of MAGs which contained contigs with laccase-coding genes.

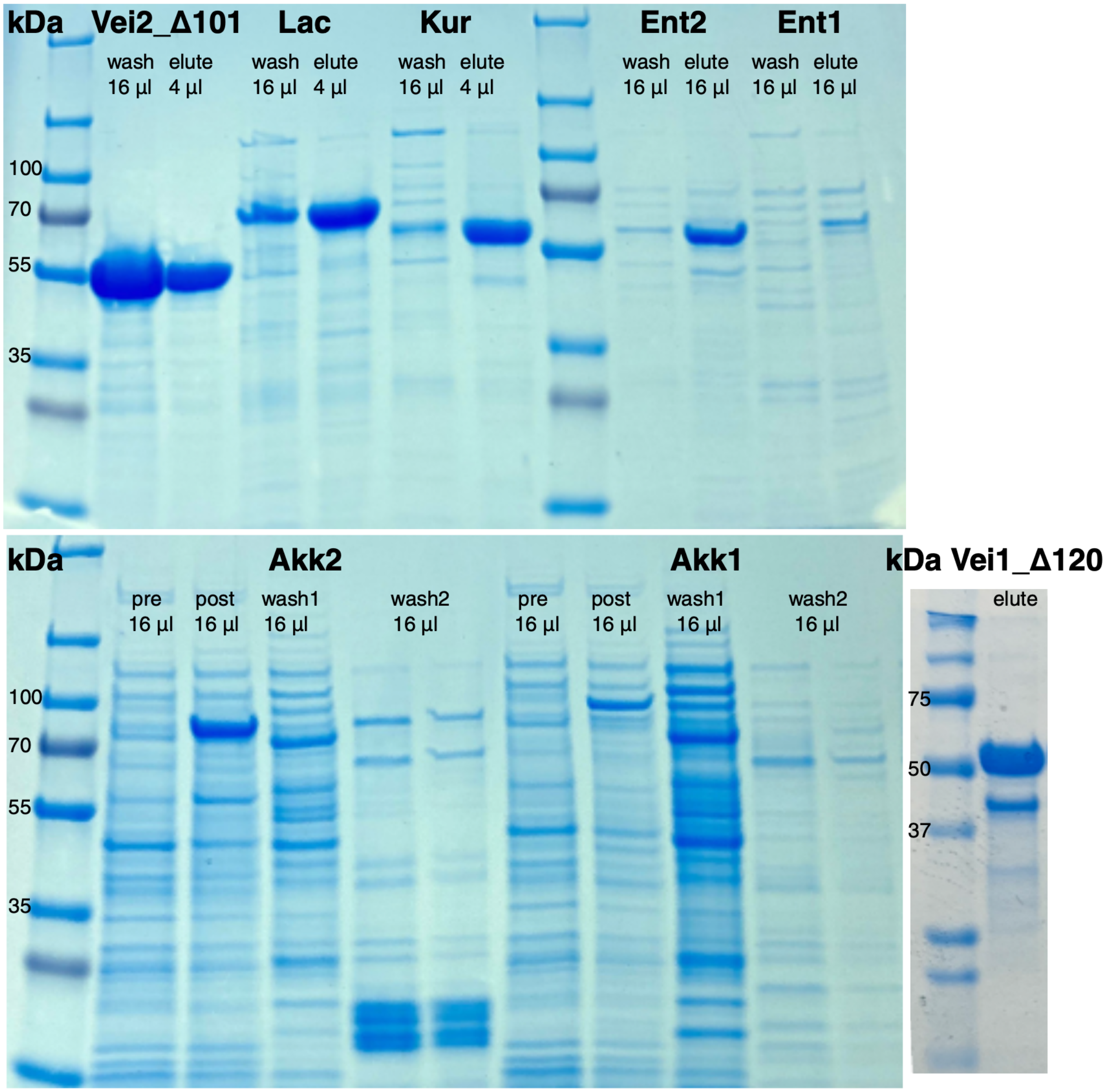


**Figure S6:** SDS-PAGE of each laccase his-column purification. Wash represents fractions from the his-column before sample elution with imidazol. Elute represents fractions after his-column elution and buffer exchange column. Vei1_Δ120, Vei2_Δ101, Lac, Kur, and Ent2 all showed reasonable purity and yield. Ent1 did not show a discernable band at the predicted size. Akk1 and 2 both showed induction by IPTG (pre- and post-induction fractions labeled accordingly), but no clear band was discernable in the final elution fractions among the significant impurities.

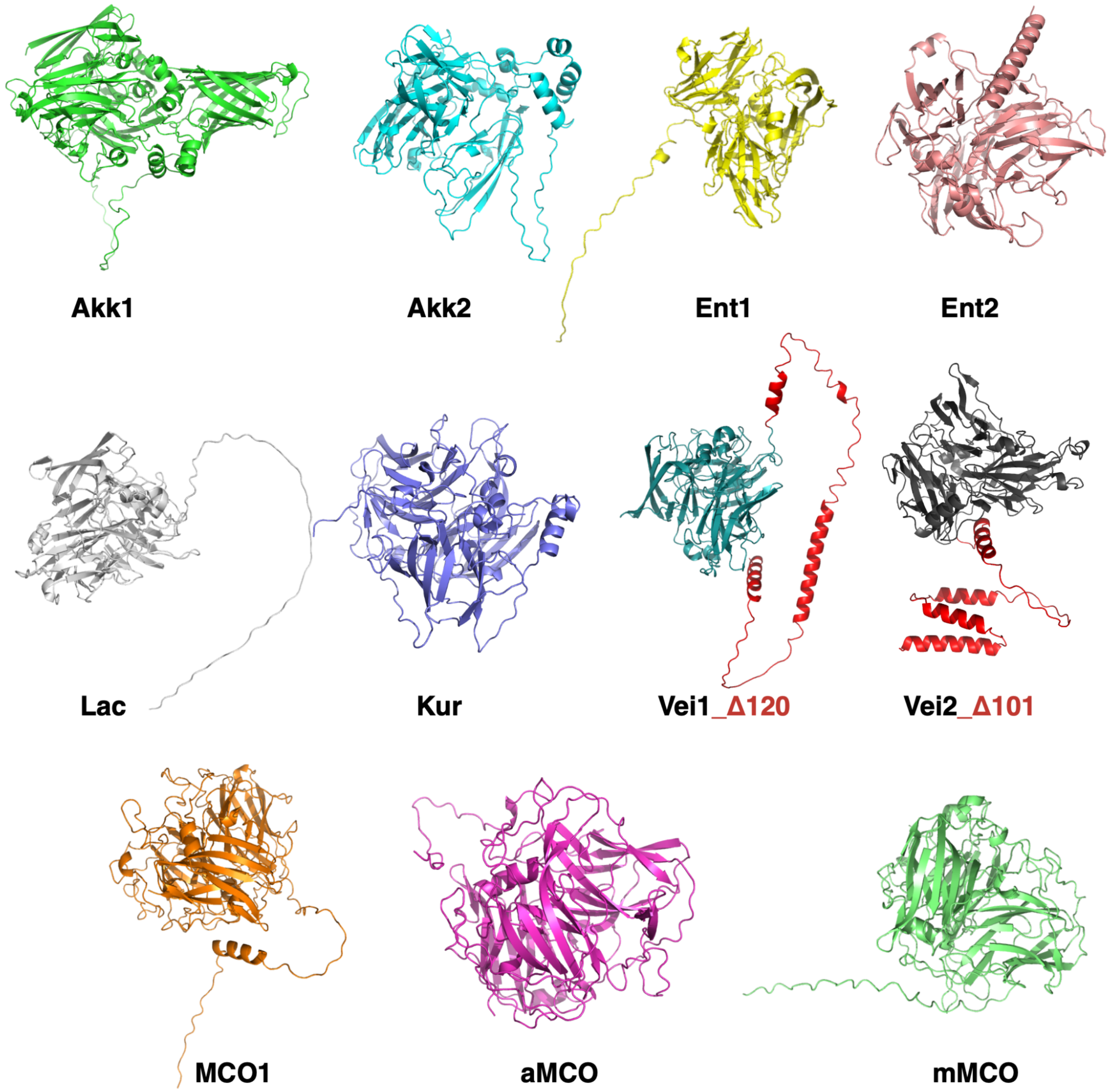


**Figure S7:** Alphafold predictions of laccase structures. For Vei1 and Vei2, the truncated portions in Vei1_Δ120 and Vei1_Δ101, respectively, are shown in red.
